# Evaluation of a BE-inactivated whole virus preparation using an encephalomyocarditis virus strain that was isolated from fatal infection in orangutans

**DOI:** 10.1101/2023.07.29.551122

**Authors:** Cathlyn Tong, Boon-Huan Tan, Richard Sugrue

## Abstract

A novel Encephalomyocarditis virus (EMCV) of the group 3 cluster (EMCV-3) was first reported in 2002 to be responsible for the deaths of orang-utans in an outbreak in the Singapore Zoo. After this first outbreak, sporadic infections among the primate population caused by EMCV-3 continued to be reported, suggesting that the virus remains prevalent in Singapore. To prevent future infections, we constructed an experimental vaccine using binary ethylenimine (BE) to inactivate the EMCV-3 virus. The immunological performance of the BE-inactivated (BEI) virus was analysed in mice and the neutralising titre of the immune sera measured against the wild-type EMCV-3. The BEI virus showed a strong immunological response in BALB/c mice at 1: 40,960 titre, suggesting that it can be used as a promising experimental vaccine candidate to prevent EMCV-3 infections.

## Introduction

Encephalomyocarditis virus (EMCV) is a non-enveloped virus of approximately 30 nm in diameter, and it belongs to the Cardiovirus A genus within the *Picornaviridae* family (Zell et al., 2017). The virus genome is a single-stranded positive-sense RNA of approximately 7.8 kb in length, which is flanked by the 5’ and 3’ untranslational regions (UTR). The virus genome encodes a leader (L) sequence and 3 polypeptides (VP1 to VP3) in the sequence of 5’-UTR-poly (C) - L - (1A-1B-1C-1D) – (2A/2B-2C/3A-3B-3C-3D) - 3’ UTR-poly (A). Based on genomic sequence analysis EMCV strains can be classified into 4 main groups, defined as EMCV-1, 2, 3 and 4 (Vyshemirskii et al., 2018). The EMCV-1 group contains 7 lineages from A to G, with the historical EMCV strains isolated from Asia, Europe, and USA. The remaining EMCV-2 group contains EMCV from Germany; with EMCV-3 group containing viruses that circulate in China and Singapore; and EMCV-4 group containing viruses from Vietnam. The virus was first identified in 1945 in a male gibbon in Miami, Florida, and since then, EMCV infections are now documented worldwide in primate and non-primate species, and in domesticated and wild animals (Lee GKW, 2015). EMCV has also been reported in humans with either asymptomatic or mild symptoms (Oberste et al., 2009), suggesting that the virus is involved in zoonotic episodes. Rats and bats found in various locations have been implicated as the virus reservoir (Doysabas et al., 2019; Liu et al., 2017).

We previously reported an outbreak of EMCV infections in orang-utans in the Singapore zoo caused by a novel EMCV-3 in 2002 (Yeo et al., 2013). The outbreak was caused by a novel EMCV strain designated SING-M105, and was associated with fatalities of the orang-utan population. Since the first outbreak reported in Singapore, EMCV continues to be sporadically detected (Lee et al., 2015), suggesting that the virus persists in the natural environment. There is an urgent need to implement effective control strategies for EMCV and one of these strategies could be the use of an effective vaccine to protect susceptible animals from EMCV infection. The persistence of this virus in the natural environment is therefore the motivation behind the current study that was to develop a specific vaccine candidate with the Singapore EMCV-3 strain. Given that the local circulating EMCV strains are likely to have specific immunogenic properties that are regional-specific, the preparations of vaccine candidates using local circulating EMCV strains is desirable. In this current study we therefore describe the development of an inactivated vaccine candidate using the EMCV-3 strain SING-M105 that was isolated from infected orang-utans during the first outbreak reported in 2002.

## Methods

### Viruses and Cells

The virus used in this study was the EMCV strain SING-M105 isolated from an orang-utan in 2002 (Yeo et al., 2013). Virus infection was carried out with African green monkey kidney epithelial (Vero E6) cells (CCL-81), obtained from American Type Culture Collection (ATCC, USA) as described in (Yeo et al., 2013). Cells were maintained in a 37°C in a humidified chamber with 5% CO_2_ in cell growth medium (DMEM supplemented with 10% heat-inactivated FCS and penicillin/streptomycin). Virus infection was performed using the EMCV-3 strain SING-M105 in maintenance medium (DMEM supplemented with 2% heat inactivated FCS and penicillin/streptomycin) at 37°C in a humidified chamber with 5% CO_2._

### Virus inactivation with binary ethylenimine (BE)

The EMCV was inactivated using binary ethylenimine (BE). 0.1M BE was freshly prepared from bromoethylamine hydrobromide (Sigma-Aldrich, USA) powder in 0.175N sodium hydroxide solution and incubated at 37°C for 1 hr as described previously (Sarkar et al., 2017). The BE solution was added to EMCV strain SING-M105 at the titre of log TCID_50_ ≥ 9 to obtain a final BE concentration of 1.5 mM, and the mixture incubated overnight at 37°C. 1M sodium thiosulphate solution was added at 10% of the volume of BE used, for 30 min at room temperature to neutralize any remaining BE in solution.

### Infectivity assay to confirm the BE inactivation

Vero E6 cells were infected with untreated EMCV and BEI-EMCV over 4 serial virus passages. Each passage (P) was prepared by infecting cells with either EMCV or BEI-EMCV and at 48 hpi, the cells and tissue culture media were harvested together. We used the cell cytopathic effect (CPE) in the EMCV-infected cells as the cue for harvesting the BEI-EMCV infected-cells. Each passage was used to infect cells and the process repeated four times in total to give P1 to P4 preparations. For each passage (P1 to P4), the untreated EMCV and BEI-EMCV preparation were serially diluted (10^-1^ to 10^-10^) and the preparations from 10^-3^ to 10^-6^ used to inoculate Vero E6 cells seeded in 96-well plate. After 72 hrs post infection (hpi) the cell monolayers were stained with crystal-violet. The presence of crystal violet staining is indicated by the dark wells and the clear wells indicate absence of cells due to cell deaths in the cell monolayer caused by virus infection.

### Immunofluorescence microscopy

The cells were fixed using 4% (w/v) paraformaldehyde in PBS at 25^°C^ for 20 mins and then washed with PBS. The cells were permeabilized using 0.1% (v/v) triton X100 in PBS at 4°C for 15 mins. The cells were washed in PBS and stained using the primary antibody and secondary antibody (conjugated to Alexa 488). The incubation time for each antibody was for 1 hr at RT. The cells were mounted in citifluor and visualized using a Nikon eclipse 80i fluorescence microscope with appropriate machine settings.

### Virus Micro Neutralization Assay

The EMCV-immune serum and pre-immune serum (control serum) was serially diluted 2-fold up to 1:1280 in maintenance medium. The serum was added to each well of a 96-well microplates in duplicate with 1 additional well as a control to monitor serum toxicity.100 TCID_50_ of EMCV strain SING-M105 was added to each well except for the serum toxicity control wells and the plate incubated at 37°C for 1hr. 1 to 2x10_4_ of Vero E6 cells was next added per well and the cells incubated at 37°C in a humidified chamber with 5% CO_2._ After 48 hrs post-infection the wells were stained using crystal-violet (prepared in formaldehyde). The absence (blue-stained cell; no CPE) or presence (clear wells, CPE) of CPE was then recorded.

### Mice immunization

1:1 (v/v) mixture of BEI-EMCV strain SING-M105 or recombinant VP1 protein prepared according or BEI-EMCV with Complete Freund’s adjuvant (Sigma-Aldrich, USA) was injected intramuscularly into five BALB/c mice at the concentration of 50 μg of antigen per mouse. The antigen concentration was determined by Bradford’s assay (Bio-Rad, USA). The mice were pre-bled before immunization and boosted 2 times, at 15 days and 36 days with 1:1 (v/v) mixture of antigen with incomplete Freund’s adjuvant (Sigma-Aldrich, USA). After the first immunization, serum samples were collected on 25 days, 46 days and 60 days and analysed for neutralising antibodies. The mice immunization procedures were conducted according to the approval from Institutional Animal Care and Use Committee (IACUC) (protocol number IACUC/16/173).

### Immunoblot Analysis

Recombinant VP1 protein and lysate prepared from Vero E6 cells infected with EMCV strain SING-M105 were separated by SDS-PAGE (Bio-Rad, USA) and the proteins transferred onto polyvinylidene fluoride membrane (Immobilion-P, Millipore, USA) according to manufacturers’ instructions. Protein bands were visualised using the ECL system (GE Healthcare, USA) and molecular masses estimated using the

√Kaleidoscope markers (Bio-Rad, USA).

### Enzyme-linked Immunosorbent Assay (ELISA)

The recombinant VP1 antigen (2 μg/ml) was used to coat the 96-well microplates overnight in binding buffer (100 mM Tris-HCL, 1 mM EDTA, 150 mM NaCl, pH8.0). The antigen-coated plates were blocked with 1% Bovine Serum Albumin for 1 hr, incubated with mouse serum for another 1 hr followed by secondary antibody, anti-mouse IgG conjugated with horse radish peroxidase (HRP), at 1:5000 dilution. Plates were washed in between incubations with PBS. After the final wash, the reaction was developed with Tetramethylbenzidine (TMB) substrate for 20 mins in the dark, stopped with 2M sulfuric acid, and absorbance measured at 450 nm using a plate reader. All measurements were conducted in triplicates.

## Results and discussion

An EMCV vaccine has been previously described for use in Australia (McLelland et al., 2005) that was inactivated using β-propiolactone (BPL). Although the trial on ungulates and chimpanzees in Australia was a success, it is not clear that this vaccine preparation would elicit the same protective antibodies against the Singapore EMCV-3 strain. BPL is currently used in several inactivated virus vaccine candidates; however BPL is also a suspected carcinogen (Information., 2022; Španinger & Bren, 2020), which may limit its general use in Singapore. In this context, BE has been used for the inactivation of various veterinary viruses to prevent livestock diseases, and we therefore evaluated BE inactivation of EMCV-3 isolate in our study. Promising virus vaccine candidates have been prepared using BE and described for use to protect livestock against Foot and Mouth disease in China (Cao et al., 2021), Bangladesh (Al Amin et al., 2020) and India (Sarkar et al., 2017). BE-inactivated vaccine candidate was reported to induce the longest duration of protective IgG levels against transmissible gastroenteritis disease in piglets when compared with vaccine candidates inactivated with formaldehyde and BPL (Zhao et al., 2020). Other examples where BEI vaccines were demonstrated to confer protection include Newcastle disease in the poultry industry (Adi et al., 2019; Aljumaili et al., 2020), Bluetongue virus infection in sheep (Bitew et al., 2019), rabies (Astawa et al., 2018) and Seneca valley virus infection in pigs (Yang et al., 2018).

We first examined if BE-treatment could be used to chemically inactivate the EMCV. Wildtype EMCV strain SING-M105 was treated with BE as described to produce the BE-inactivated EMCV strain SING-M105 isolate (BEI-EMCV). The efficiency of BE-inactivation was determined by serially passaging the BEI-EMCV four times in Vero E6 cells. The inactivation of the BEI-EMCV was confirmed for each of the four serial passages by examining cytotoxicity in Vero E6 cells (**Figure 1A**). The infectious EMCV and BEI-EMCV preparations were serially diluted in the range 10^-1^ to 10^-10^ and the 10^-3^ to 10^-10^ dilutions were used to infect the Vero cells. These dilutions were used to avoid the cell effects related to host cell factors / antivirus effectors that are expected to be released into the tissue culture media as a result of CPE induced in the EMCV-infected cells at 48 hpi. After 72 hpi the cells were stained using crystal violet and imaged using light microscopy. The four serial passages of BEI-EMCV showed no signs of infectivity, while in the EMCV-infected cells, cell death at between the 10^-5^ and 10^-8^ virus dilutions was noted. At the end of the 4^th^ passage Vero E6 cells were infected with untreated EMCV and BEI EMCV, and after 18 hpi the cells were co-stained using anti-VP1 and Evans Blue (EB), and examined by immunofluorescence (IF) microscopy (**Figure 1B**). The anti-VP1 is a mouse polyclonal antibody that was produced using the recombinant expressed VP1 protein of the EMCV-3 SING 105 strain (**SFigure 1**). While EB-staining allows imaging of all cells in the field of view, anti-VP1 staining was observed in EMCV-infected cells. No anti-VP1 staining was observed in either the mock-infected or BEI-EMCV-infected cells, confirming the inactivation of EMCV strain SING-M105. Collectively, the cytotoxicity and IF microscopy data demonstrated that BE-treatment can successfully inactivate EMCV strain SING-M105.

**Figure 1.**
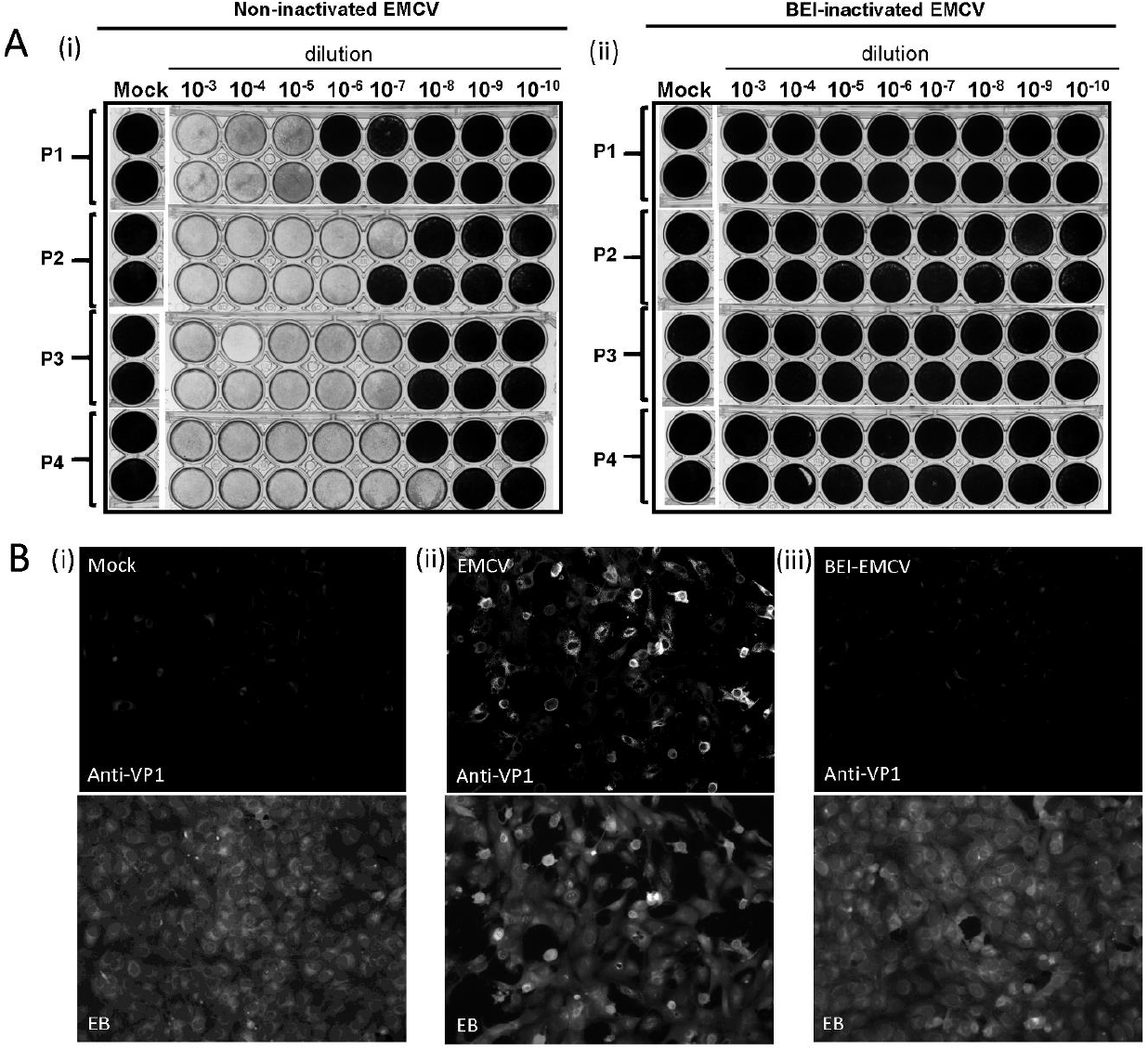
Inactivation of EMCV using treatment with binary ethylenimine (BE). **(A)** Cytotoxicity assay to measure infectivity in BE inactivated (BEI)-EMCV. Vero E6 cells were infected with untreated EMCV and BEI-EMCV over 4 serial virus passages (P1, 2, 3 & 4 represents sequential passages). Each virus preparation was serially diluted (10^-3^ to 10^-10^). (i) Untreated EMCV and (ii) BEI-EMCV dilutions was added to the respective wells (in duplicate) and after 72 hrs post infection (hpi) the cell monolayers were stained with crystal-violet. The presence of crystal violet staining is indicated by the dark wells and the clear wells indicate absence of cells due to cell deaths in the cell monolayer caused by virus infection. Also shown are wells from the mock-infected cells (mock) stained at 72 hpi. **(B)** Examination of Vero E6 cells infected with untreated EMCV and BEI-EMCV using immunofluorescence (IF) microscopy. Vero E6 cells were (i) mock-infected, (ii) infected with untreated EMCV and (iii) BEI-EMCV respectively. After 18 hrs post-infection (hpi) the cells were co-stained with anti-VP1 and Evans Blue (EB) and examined using IF microscopy (objective X 20). The cells were viewed using identical machine/camera settings.

We next examined the immunogenicity response of the BEI-EMCV in BALB/c mice as described in the methods. Five BALB/c mice were pre-bled and intra-muscularly immunised with 3 doses of the BEI-EMCV. The pre-immune serum (PIS) and immune serum (EMCV-IS) were collected from each animal and the immunogenicity data from the serum of one representative mouse is presented (**Figure 2**). We examined the immunoreactivity of PIS and the EMCV-IS by immunoblotting with EMCV-infected VERO E6 cell lysate (**Figure 2A**). A 32 kDa representing the size of VP1 protein as highlighted by the black arrow, was observed by immunoblotting using the EMCV-IS. A second protein band at about 25 kDa was also detected, which is the expected size for the EMCV-VP3 protein. The protein bands were not detected by immunoblotting with the PIS. The EMCV-infected cells were also co-stained using EB and either the PIS or EMCV-IS and examined by IF microscopy (**Figure 2B**). The EMCV-IS (Anti-EMCV) showed specific florescence staining with the EMCV-infected Vero cells while no significant staining was observed in PIS-stained cells. These data provide evidence that the EMCV-IS was able to recognise the EMCV capsid protein.

**Figure 2.**
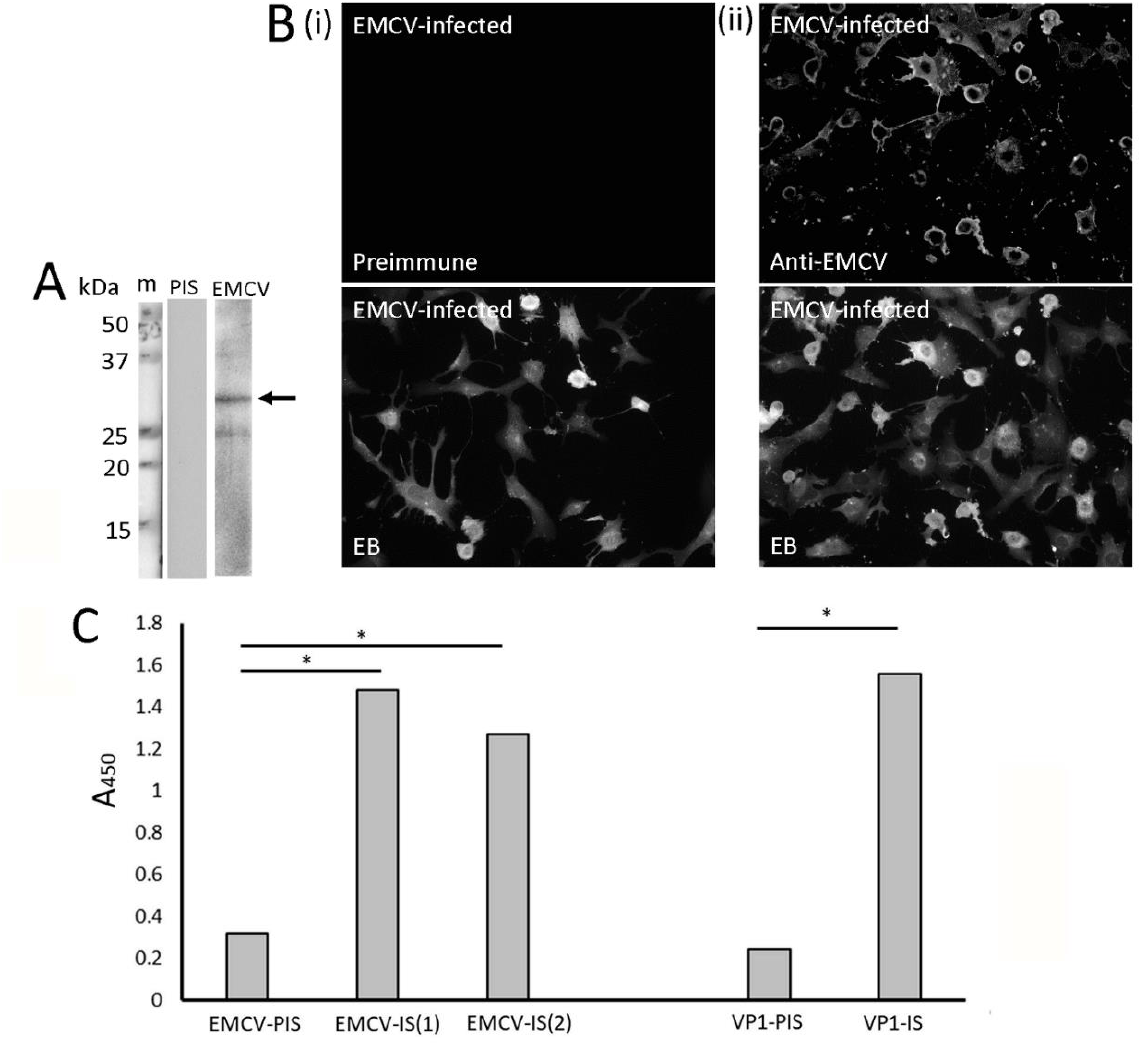
The immunogenicity of the binary ethylenimine inactivated (BEI)-EMCV immune serum. Five BALB/c mice were pre-bled and intra-muscularly immunised with 3 doses of the BEI-EMCV. **(A)** Immunoblot analysis of pre-immune serum (PIS) and EMCV-immune serum (EMCV) in EMCV-infected cells. The PIS and EMCV were immunoblotted with the cell lysate from EMCV-infected Vero cells. The arrow highlights the position of VP1 protein at 32 kDa size. **(B)** Vero E6 cells were infected with EMCV and the infected cells allowed to react with pre-immune serum (Preimmune) and immune serum (anti-EMCV) and examined with immunofluorescence microscopy (objective X 20) The cells were viewed using identical machine/camera settings. Cells were co-stained using Evans blue (EB). **(C)** The immune-reactivity of EMCV-IS was examined by ELISA using recombinant VP1 protein-coated plates. VP1 recombinant protein were coated at 1:5000 dilution, and allowed to react with pre-immune (EMCV-PIS) and immune sera raised against BEI-EMCV defined as 2^nd^ bleed [EMCV-IS (1)] and final bleed [EMCV-IS (2)] respectively. Also shown is the reactivity using VP1-PIS and VP1-IS. The immune-reactivity was measured at A=450nm. Average readings of triplicate results are given. The p value is <0.01 and was performed with the paired sample t-test.

The recognition of the VP1 protein by the EMCV-IS was also examined by ELISA using the recombinant expressed VP1 protein to coat the ELISA plates (**Figure 2C**). In both the 2nd bleed (harvested at 45 days) and the final bleed (harvested at 60 days) EMCV-IS were immune-reactive with the VP1 antigen. When the serum collected at the final bleed was evaluated, reactivity to the VP1 antigen at 1:5000 dilution was observed with similar magnitude of reactivity to that of second bleed. The pre-immune serum only showed low levels of immune-reactivity with the VP1 antigen, which is presumably due to background signal in the assay. As a control we compared the pre-immune serum and the immune serum obtained from mice inoculated with a purified VP1 antigen (VP1-IS), which showed similar immuno-reactivity of EMCV-IS and VP1-IS, and low baseline signal using the pre-immune serum from mice inoculated with the recombinant VP1 protein.

These data indicated that the BEI-EMCV was capable of inducing an immune response in the mice and we next examined if the BEI-EMCV preparation was able to produce a neutralising antibody response that were capable of blocking EMCV infection. This assay was performed using a micro-neutralisation assay in which the wildtype EMCV strain SING-M105 was pre-incubated with either the PIS or EMCV-IS obtained from each of the five mice before infecting the Vero E6 cells (**Figure 3**; represented as Pre-immune Serum and EMCV Immune Serum respectively). The serum harvested from each mouse was serially diluted (2-fold) from 1:10 up to a dilution of 1:82,190. As a control the PIS from each mouse was also harvested and a similar 2-fold serial dilution performed. The EMCV was pre-incubated with either the PIS or the EMCV-IS dilutions from each of the five mice and the virus then used to infect Vero E6 cells. The cells were incubated for 72 hrs after which the cell monolayers were stained using crystal violet to detect cytotoxicity. In the absence of

**Figure 3.**
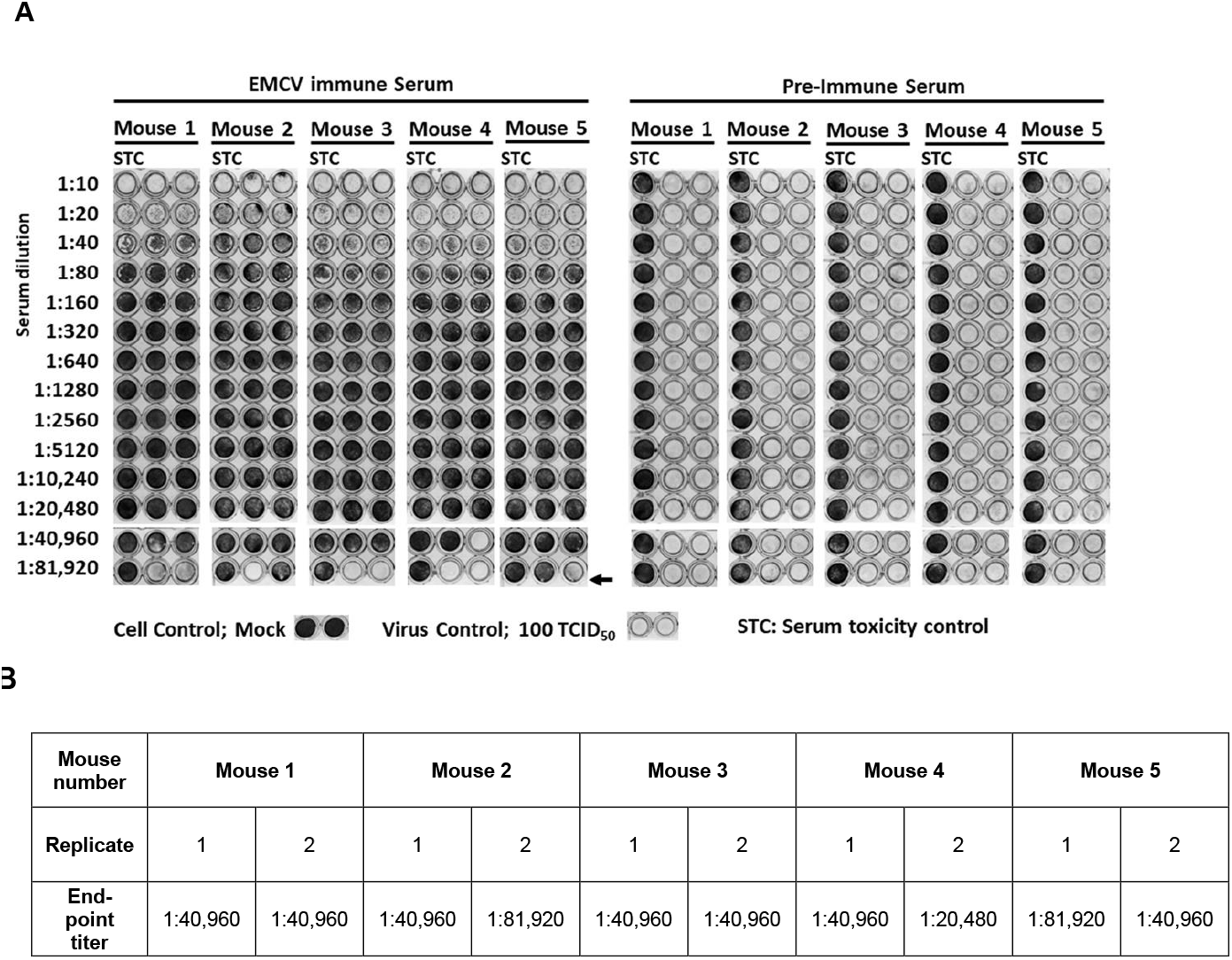
Micro-neutralisation analysis of immune serum (anti-EMCV) raised against BEI-EMCV. Five BALB/c mice were pre-bled and intra-muscularly immunised with 3 doses of the BEI-EMCV. The pre-immune and immune sera were collected from each of the five mice and analysed. Wild-type EMCV strain SING-M105 was added at 100 TCID_50_ per well in the presence of pre-immune and immune-serum for mouse 1 to 5. The serum from each mouse was diluted serially two-fold from 1:10 to 1:81290. After 72 hrs post-infection the Vero E6 cells were stained with crystal-violet to detect the presence of cytotoxicity. **(A)** Micro-neutralisation of EMCV immune and pre-immune sera in microplate. The clear wells indicated cytotoxicity (absence of cells) and the dark wells indicate the absence of cell cytotoxicity (stained cells). STC represents serum toxicity control. The black arrow indicates the end point in the titration. **(B)** A summary of the end-point titre of the micro-neutralisation assay from each of the five mice. Replicates 1 and 2 represent results conducted in duplicates.

cytotoxicity the crystal violet stains the cell monolayer and can be distinguished from the well exhibiting cytotoxicity (clear wells). The endpoint in the assay was taken when at least one of the two replicates for each dilution showed CPE. On this basis we estimated that in mice 1 to 5, an end point titration of 1:40,960 was noted for the EMCV-IS. There was no virus neutralisation observed in the virus incubated with the PIS as all cells were infected. Serum toxicity on the Vero E6 cells was also performed in parallel with the micro-neutralisation but in the absence of EMCV strain SING-M105. Although the BEI-EMCV preparation was well-tolerated in the inoculated mice with no unusual adverse effects, up to a dilution of 1:40 the EMCV-IS appeared to show serum toxicity on the Vero cells when compared with the PIS. However, no toxicity on Vero E6 cells were observed at EMCV-IS dilutions greater than 1:40. Since the induction of cytokines and other factors facilitate humoral immune responses this apparent cytotoxicity at low serum dilutions may be an indirect indicator of a robust immune response in these animals. The results from the micro-neutralisation assay showed that the BEI-EMCV was able to produce high neutralising antibodies to the EMCV strain SING-M105. This finding was in contrast to our earlier investigation where we used recombinant SING-M105 VP1 protein as an antigen to immunize Balb / C mice. The immune serum of mice immunised using the recombinant VP1 was performed using micro-neutralisation assay method described above. This analysis indicated that although the recombinant VP1 was able to elicit a neutralising antibody response in an immunised mouse, the neutralising antibody responses recorded was significantly lower than in mice immunised with the BEI-EMCV (**SFigure 2**). Interestingly, a panel of monoclonal antibodies was generated from the mouse immunised with the recombinant VP1, and one of these antibodies (MAb 5C2) exhibited virus neutralising activity that was higher than recorded for the VP1 immune serum (**SFigure 3**). Although the neutralising titre for the VP1 monoclonal antibody was lower than that recorded for the serum obtained from the BEI-EMCV immunised animals, these data suggest the possibility of using therapeutic antibodies for passive immunisation as an option for the rapid treatment of EMCV-infected animals.

## Conclusions

EMCV-3 continues to be present in the natural environment, suggesting the potential risk of a disease outbreak at short notice in orang-utans held in captivity. Measures are needed to be in place to mitigate the infection risk and this would include continuous surveillance in animals that are susceptible to risk of fatal infection. This includes orang-utans as well as animals that constitute a natural reservoir for the virus e.g. rodents. The development of an EMCV vaccine against circulating Singapore EMCV strains represents an important factor in the control of EMCV infection. The BEI-virus vaccine described in this study was able to induce high neutralising antibody titres in immunised mice and it was well tolerated in these immunised animals. This suggests that the BEI vaccine candidate could be used in future to mitigate future EMCV-3 outbreaks in Singapore and in the region. There are also several advantages of using BE to inactivate viruses. Vaccination with BEI vaccines are known to induce long-lasting protective antibodies with cellular immunity effects (Zhao et al., 2020) and BEI vaccines rapidly achieved peak antibody titres within a short span of vaccination (Al Amin et al., 2020). This suggests that upon vaccination, protection can be rapidly achieved within a short immunisation time, and would be useful in employing ring-vaccination as part of disease management before the infection risk escalates into a potential outbreak.

Our current finding has shown that the BEI-EMCV is able to produce a strong neutralisation titre, and our future studies will demonstrate this protective response in a suitable animal model. Past reports have successfully described the susceptibility of both black 6 (Ohtaki et al., 2012) and Balb / C mice (Zhu et al., 2011 and 2015) to EMCV infections and using these models to study disease pathogenicity. We would anticipate using either mice model to develop challenge models with the Singapore EMCV strain. Since protecting orang-utans from EMCV infection was the primary motivation for this study, future studies will also examine the effectiveness of the BEI-EMCV-preparation and a suitable adjuvant in producing a protective immune response in orang-utans.

## Authors contributions

Conceptualisation, BHT, RJS; Methodology, CT; Data analysis, CT; Data curation, BHT, RJS; writing, reviewing, editing BHT, RJS; project administration, BHT, RJS.

## Ethical Approval

The mice immunization procedures were conducted according to the approved protocols from Institutional Animal Care and Use Committee (IACUC) (protocol number IACUC/16/173) from DSO National Laboratories.

## Consent for publication

All authors consented to the publication.

## Acknowledgements

We thank Koon-Hui Lee, Mok-Wei Heng, Ching-Ging Ng, and Teck-Choon Ayi from DSO National Laboratories for their technical assistance. We also thank Dr Peter Kirkland (Elizabeth Macarthur Agriculture Institute, Australia) for his technical advice offered during the course of this study.

