## Supplementary Figures for "Evaluation of a BE-inactivated whole virus preparation using an encephalomyocarditis virus strain that was isolated from fatal infection in orangutans"

**Supplementary Figure 1.**

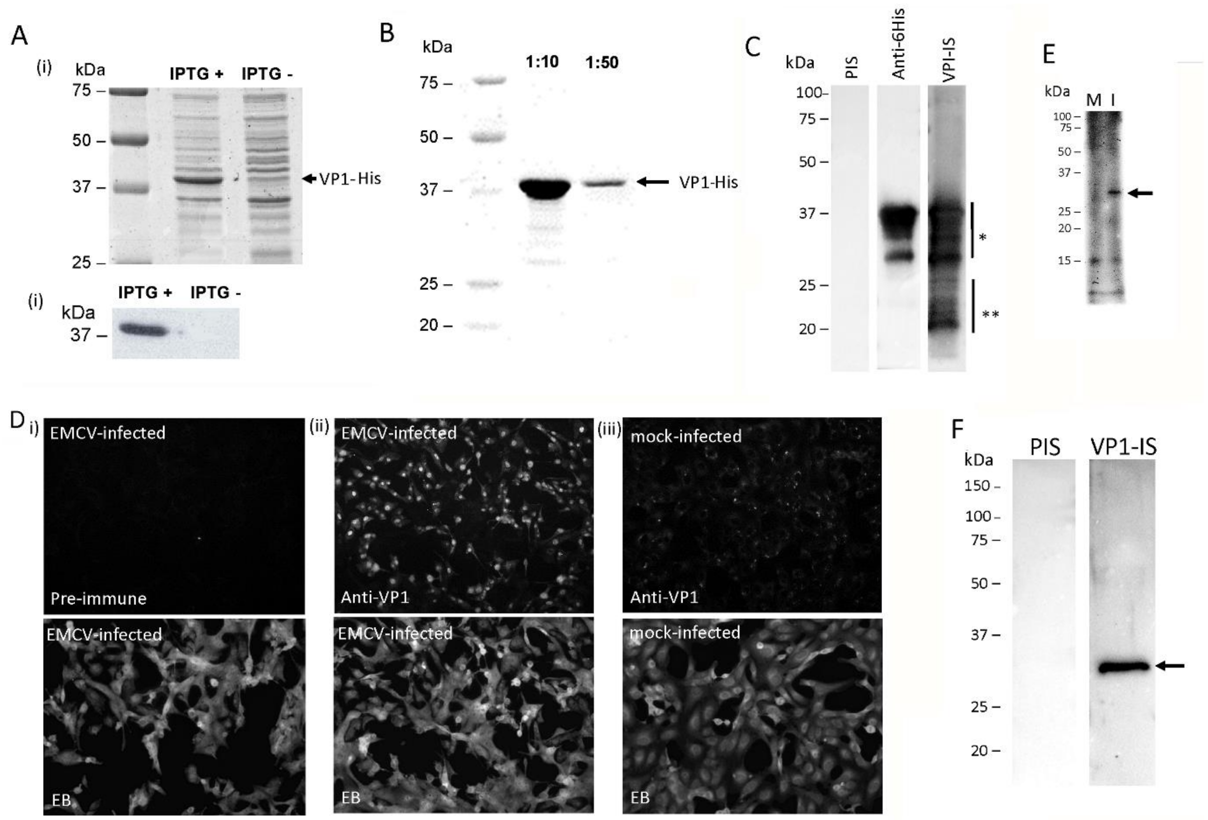


**SFigure 1.** Production of anti-VP1 antibody using recombinant expressed VP1 protein. Recombinant VP1 protein was expressed in *E. coli* BL21(Des)pLysS cells and the crude lysate was shown in **(A)** **(i)** Coomassie-stained SDS-PAGE gel and **(ii)** immunoblot analysis with IPTG-induced (IPTG +) and uninduced (IPTG -) condition. The arrow highlighted the position of VP1-6His protein at 38 kDa size. **(B)** Immunoblot analysis of purified VP1-6His reacting with anti-6-His of VP1 protein at 1:10 and 1:50 dilution. The arrow highlighted the position of VP1-6His protein at 38 kDa size. **(C)** Immunoblot analysis of the VP1-6His protein reacting with pre-immune serum (PIS), anti-6His and immune serum (VPI-IS) to recombinant VP1 protein; * and ** represents different protein species of degraded recombinant VP1 protein. **(D)** Immunofluorescence microscopy of Vero E6 cells infected with EMCV strain SING-M105 and stained with **(i)** pre-immune and **(ii)** immune serum (anti-VP1) collected from one representative mouse. **(iii)**Mock-infected cells were also stained with anti-VP1 serum. Infected cells were examined using immunofluorescence microscopy (objective X 20). **(E) & (F)** Immunoblot analysis of the pre-immune (PIS) and immune (VP1-IS) sera against **(E)** mock-infected (M) and wild-type EMCV (I) strain SING-M105-infected (I) Vero E6 cell lysate and **(F)** purified wild-type EMCV strain SING-M105 protein separated on the SDS-PAGE gel. The black arrow highlighted the native VP1 viral protein banding at 32 kDa size in both figures.

**Supplementary Figure 2.**

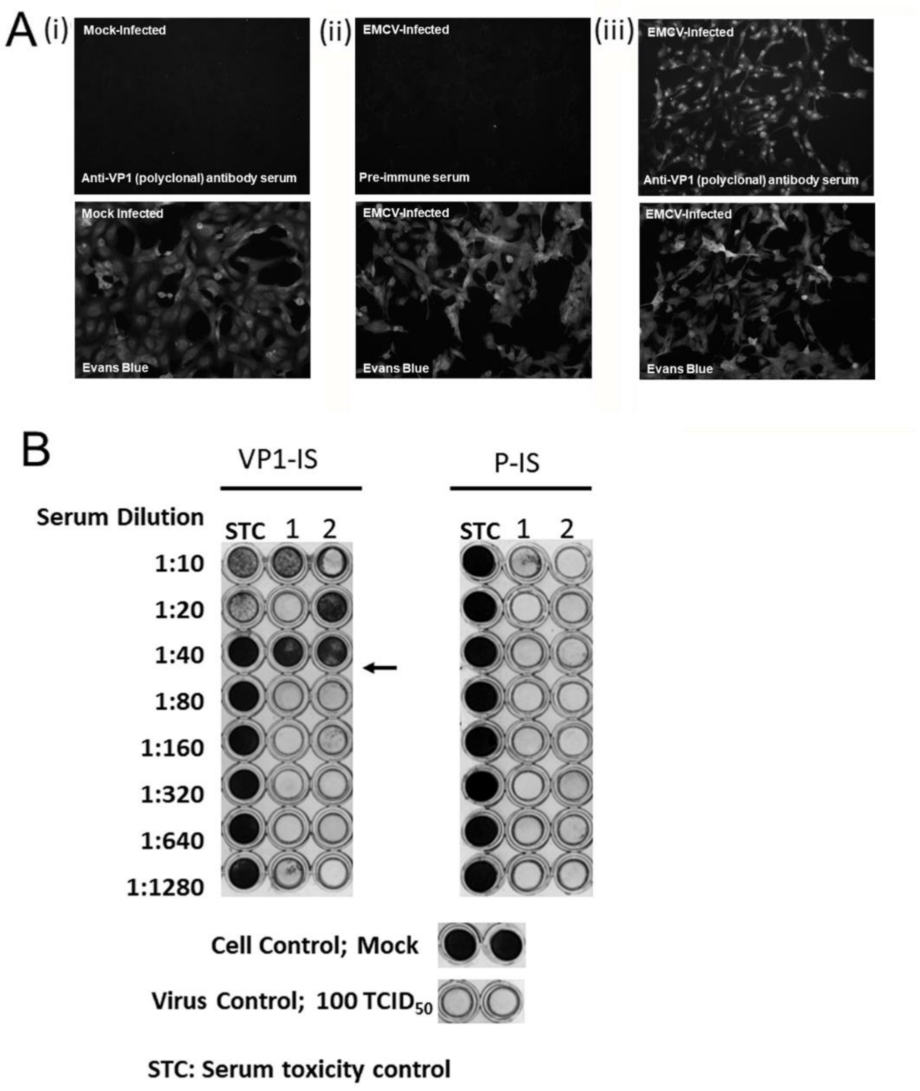


**SFigure 2.** The immunogenicity of immune serum raised against recombinant VP1 protein. The pre-immune and immune sera collected from one representative immunised mouse is analysed. **(A)** Immunofluorescence microscopy of Vero E6 cells **(i)** mock-infected and infected with EMCV strain SING-M105, and stained with **(ii)** pre-immune and **(iii)** immune serum (polyclonal). Infected cells were examined using immunofluorescence microscopy (objective X 20). **(B)** Micro-neutralisation analysis of immune serum raised against recombinant VP1 protein. One BALB/c mouse was pre-bled and intra-muscularly immunised with 3 doses of the recombinant VP1 protein. The pre-immune (P-IS) and immune sera (VP1-IS) were collected. Wild-type EMCV strain SING-M105 was added at 100 $TCID_{50}$ per well in the presence of pre-immune and immune-serum. The serum from each mouse was diluted serially two-fold from 1:10 to 1:1280. After 72 hrs post-infection the Vero E6 cells were stained with crystal-violet to detect the presence of cytotoxicity. The clear wells indicated cytotoxicity (absence of cells) and the dark wells indicate the absence of cell cytotoxicity (stained cells). STC represents serum toxicity control. The black arrow indicates the end point in the titration.

**Supplementary Figure 3.**

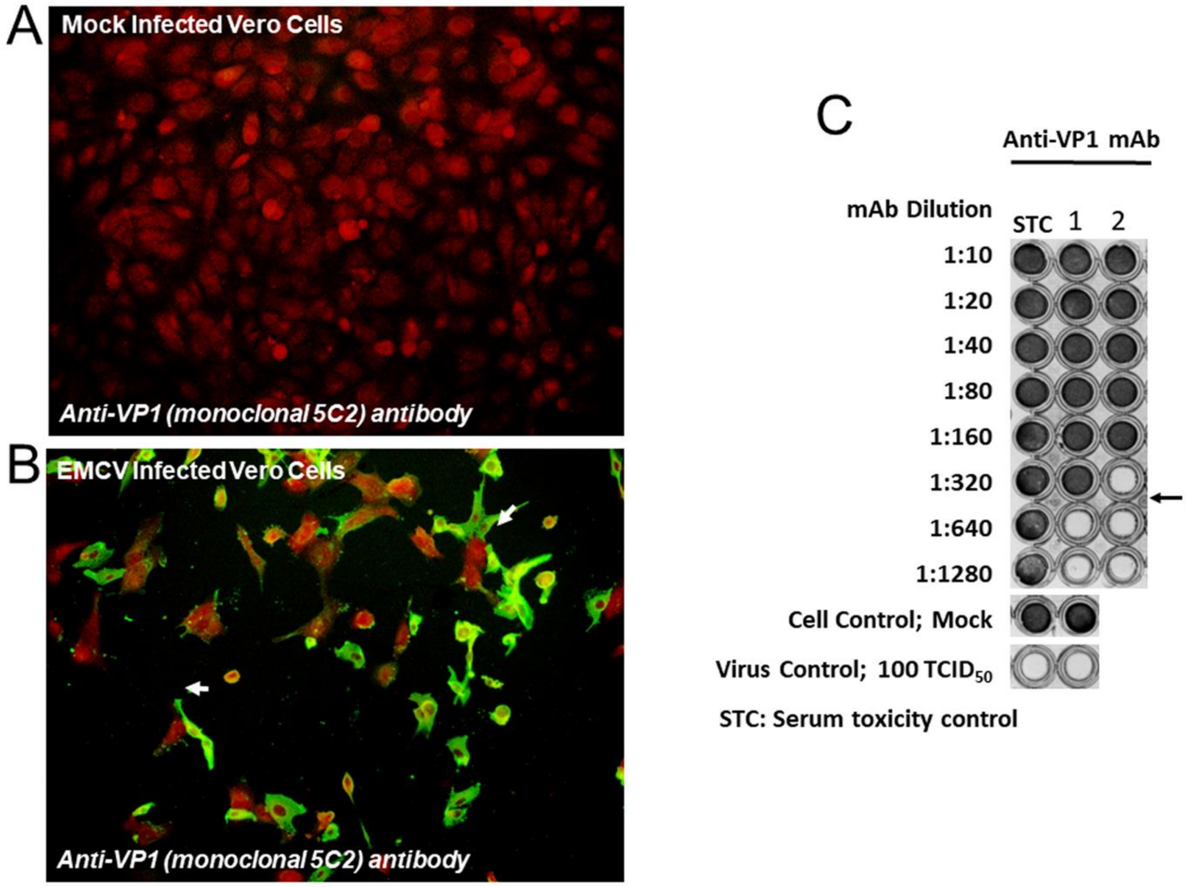


**SFigure 3.** The immunogenicity of monoclonal antibody raised against recombinant VP1 protein. The monoclonal antibody (MAb) was generated from hybridoma using standard methodology. Immunofluorescence microscopy of Vero E6 cells **(A)** mock-infected and **(B)** infected with EMCV strain SING-M105, and stained with anti-VP1 **(**monoclonal 5C2). Cells were co-stained with Evans Blue and examined using immunofluorescence microscopy (objective X 20). The reactivity of the VP1 MAb was shown as green florescence and in its absence, the uninfected cells stained red with Evans Blue. **(C)** Micro-neutralisation analysis of monoclonal antibodies (anti-VP1 MAb) raised against recombinant VP1 protein. Wild-type EMCV strain SING-M105 was added at 100 $TCID_{50}$ per well in the presence of MAb. The MAb was diluted serially two-fold from 1:10 to 1:1280 and in duplicate. After 72 hrs post-infection the Vero E6 cells were stained with crystal-violet to detect the presence of cytotoxicity. The clear wells indicated cytotoxicity (absence of cells) and the dark wells indicate the absence of cell cytotoxicity (stained cells). STC represents serum toxicity control. The black arrow indicates the end point in the titration.
